# Enhanced retinoid response by a combination of the vitamin A ester retinyl propionate with niacinamide and a flavonoid containing *Ceratonia siliqua* extract in retinoid responsive *in vitro* models

**DOI:** 10.1101/2020.09.26.315093

**Authors:** Eric C.S. Lam, Rui Li, MyriamRubecca Rodrigues, Laura Vires, Rachel L. Adams, Joseph D. Sherrill, John E. Oblong

**Affiliations:** Procter and Gamble International Operations SA SG Branch, Singapore, Singapore; The Procter and Gamble Company, Cincinnati, OH, USA

**Keywords:** retinyl propionate, keratinocytes, niacinamide, *Ceratonia siliqua*

## Abstract

**OBJECTIVES:** Retinoids have been used for decades as efficacious topical agents to treat photoaged skin. The purpose of our present research is to evaluate whether the activity of the Vitamin A ester retinyl propionate (RP) can be enhanced by niacinamide (Nam) and a flavonoid containing *Ceratonia siliqua* (*CS*) fruit extract in retinoid responsive *in vitro* models.

**METHODS:** RP was tested alone and in combination with Nam and *CS* in an RARα reporter cell line for promoter activation and compared to *trans*-retinoic acid (tRA) activation. These treatments were also tested in keratinocytes for gene expression profiling by qPCR using a panel of 40 retinoid responsive genes.

**RESULTS:** tRA or RP elicited RARα reporter activation in a dose dependent manner. The combination of 0.5 μM or 2 μM RP with 10 mM Nam had a 56% and 95% signal increase compared to RP, respectively. The addition of 1% *CS* to 0.5 μM or 2 μM RP with 10 mM Nam elicited a further increase of 114% and 156%, respectively, over RP and Nam combinations. All retinoids elicited an increase in expression of 40 retinoid sensitive genes over control levels. Of the 40 genes 27 were enhanced by either 0.5 μM RP or 2 μM RP with 10 mM Nam and 1% *CS*. Nam or *CS* had very modest activity in both models.

**CONCLUSION:** The combination of RP with Nam and *CS* showed a higher retinoid response than RP in two separate retinoid responsive *in vitro* models. We hypothesize Nam and *CS* enhances RP activity by modulating metabolism to tRA via increasing NAD^+^ pools and inhibiting reduction of retinal (RAL) back to retinol, respectively. The findings provide evidence that this combination may have enhanced efficacy for treating the appearance of photoaged skin.

## Introduction

The skin is the largest organ of the human body and one of its primary functions is to provide protection from damaging environmental stressors such as solar UV radiation, carbon emissions, and pollution. Cumulative exposure to these damaging stressors leads to premature structural and functional changes that manifest as older aged skin appearance and has been ascribed as photoaging. Solar UV radiation is considered the most significant stressor and has been estimated to account for ∼85% of the premature aging cascade in skin [1, 2].

Retinoids are lipophilic vitamin A derivatives that have been used for decades in the treatment of photoaged skin [3]. Mechanistically, retinoids play an important role in epidermal homeostasis, particularly regulating proliferation and differentiation of keratinocytes and maintaining epidermal thickness [4-6]. The primary active retinoid form is *trans*-retinoic acid (tRA), which binds to members of the retinoic acid receptor (RAR) family of nuclear hormone receptors. Upon binding, the RAR complex translocates into the nucleus to activate selective gene expression [7, 8]. While systemic vitamin A is obtained via diet and oral absorption, retinoids can also become bioavailable when applied topically [9, 10]. Epidermal keratinocytes and dermal fibroblasts are enzymatically capable of converting ROL and retinyl esters to tRA via an NAD+ dependent oxidative pathway [11-13].

RP is a vitamin A ester analogue and has been reported to clinically impact photoaged skin with minimal irritation [14-16]. In addition to its overall efficacy and skin tolerability, RP has also been reported to have a better chemical stability profile compared to other retinyl esters, thereby increasing its half-life on the skin’s surface during topical delivery [17]. In the present work we wished to further enhance efficacy potential and minimize any risk of irritation by identifying compounds which could boost RP activity in retinoid responsive models. Previous approaches to increase retinoid activity and overcome retinoid resistance have focused on inhibition of P_450_ hydroxylases, particularly members of the CYP26 family, which metabolize tRA to the inactive form of 4-oxo-retinoic acid [18]. This has led to the development of retinoic acid metabolism blocking agents (RAMBA) such as the azoles liarazole and rambazole, potent inhibitors that can reduce retinoid resistance [19]. We evaluated an alternate approach to retinoid enhancement by focusing on the metabolism pathway of RP to tRA via the ROL and RAL intermediates that require NAD^+^ as a co-factor. We hypothesized that the NAD^+^ precursor Nam would facilitate this conversion by incorporating into cellular NAD^+^ pools and there by enhance RP activity. This is of particular relevance in photoaged skin since it has been established that cellular NAD^+^ pools in skin decline significantly with age, which impacts the role it has in maintaining skin homeostasis (20, 21). Additionally, we evaluated the flavonoid content in a *Ceratonia siliqua* (carob) fruit extract (*CS*, Silab, France) since flavonoids have been reported to inhibit AKR1B10 and carbonyl reductases, enzymes that can reduce RAL back to ROL [22, 23]. *CS* sourced raw materials are of keen interest for human health benefits, including topical skin care usage, based on their flavonoid and polysaccharide content [24, 25]. A total polyphenolic Folin-Ciocalteu quantification assay was performed on the *CS* fruit extract used in this work and it was calculated to have a total flavonoid content of 104 μg/ml (data not shown, Polyphenol Quantification Assay Kit, Bioquochem, Asturias, Spain).

A RARα reporter (Luc)-HEK293 cell line that contains a stably transfected luciferase gene under control of a retinoic acid receptor response element along with a full length human RARα gene (Promega, Madison, WI) was used to quantify the effect of retinoids on induction of luciferase activity. This reporter system requires the presence of tRA and thus is an indirect measure of enzymatic conversion of ROL and RP into tRA inside the cells. Briefly, cells were grown in assay media and seeded into a 96-well plate at a density of ∼30,000 cells per well. Cells were exposed to respective treatments for 24 hours after which viability was measured using CellTiter-Fluor as per manufacturer’s instructions (Promega, Madison, WI). Subsequently, luciferase activity was measured by BioGlo reagent as per manufacturer’s instructions (Promega, Madison, WI). CellTiter and Bioglo luminescence measurements were performed using a Cytation 3 Imaging Plate Reader (BioTek Instruments, VT, USA).

Cells were treated with tRA between 0.00001 to 0.1 μM to establish a luciferase signal range (Figure 1a). All tRA doses significantly induced luciferase activity by 618% (at 0.00001 μM), 682% (at 0.0001 μM), 928% (at 0.001 μM), 1203% (at 0.01 μM), and 1895% (at 0.1 μM) as a percentage of vehicle control. Cells were also treated with 0.02, 0.1, 0.5, 2, or 10 μM RP (Figure 1b). RP significantly induced luciferase activity by 90% (*p<0.05) at 0.1 μM and by 413%, 593%, and 550% at 0.5, 2 or 10 μM, respectively (***p<0.001) as a percentage of vehicle control. 0.5 μM and 2 μM RP were selected for combination testing (Figure 1c) since the basal signal increase by RP alone was in the lower range of the tRA dose response curve (Figure 1a). When 0.5 μM or 2 μM RP was tested in combination with 10 mM Nam, there was a 56% and 95% increase in signal over RP alone, respectively. Importantly, the combination of 1% *CS* and 10 mM Nam with 0.5 μM or 2 μM RP further increased the measured luciferase signal by an additional 115% and 157%, respectively, over RP alone.

**Figure 1.**
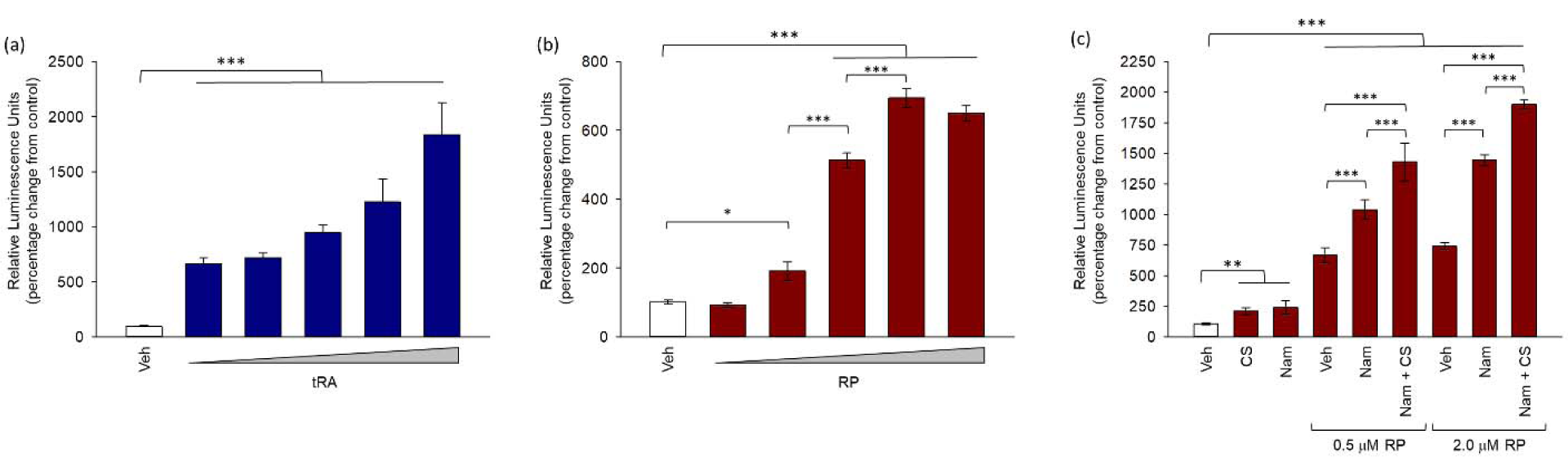
RARα activation by RP can be enhanced when combined with niacinamide and *Ceratonia siliqua*. HEK293 cells containing a stable RARα reporter construct were treated for 24 hours with tRA, ROL, RP, niacinamide (Nam), *Ceratonia siliqua* (*CS)*, or a combination of RP with Nam and *CS*. Luciferase activity was quantitated, normalized to control cells and signal compared to vehicle control treatment (Student’s t-test, \*\*\**p*<0.001, ** *p*<0.005, * *p*<0.01, n=3). (a) Normalized luciferase signal from cells treated with 0.00001, 0.0001, 0.0001, 0.01, or 0.1 μM tRA (blue bars, increasing dose represented by rising triangle) compared to vehicle control (white bar). (b) Normalized luciferase signal from cells treated with 0.02, 0.1, 0.5, 2, or 10 μM RP (red bars, increasing dose represented by rising triangle) compared to vehicle control (white bar). (c) Normalized luciferase signal from cells treated with 1% *CS*, 10 mM Nam, 0.5 μM RP, or 2.0 μM RP and the combinations of 0.5 μM RP or 2.0 μM RP with 10 mM Nam or with 10 mM Nam and 1% *CS* (red bars) compared to vehicle control (white bar).

We next evaluated the impact of RP alone and in combination with *CS* and Nam on the expression profiles of 40 retinoid responsive genes by qPCR. hTERT keratinocytes were cultured in Epilife media with full supplements and treated for 24 hr with vehicle (DMSO) or retinoid treatments. Total RNA was isolated from cell lysates, quantitated, and cDNA generated. cDNA was plated onto a Wafergen MyDesign SmartChip (TakaraBio, p/n 640036) using the Wafergen Nanodispenser. qPCR was then performed on the chip and relative expression values of the 40 target genes were normalized to the geometric mean of 4 housekeeping genes (*ACTB, B2M, GAPDH*, and *PPIA*) and fold changes over vehicle-treated cells was evaluated for significance using a Student’s t-test. A complete list of the target genes analysed can be found in Supplementary Table 1.

Treatment with 0.1 μM tRA, 0.1 μM ROL, 0.1 μM RP or 0.5 μM RP all significantly increased expression of the target genes. 0.1 μM and 0.5 μM RP were selected based on these concentrations eliciting a similar expression induction level as measured in the lower range of a tRA dose response curve (data not shown). All 40 target genes showed an increase in expression level in response to RP and 16 were RP dose sensitive (Supplementary Table 1). Nine of the dose responsive genes (ALDH1A3, ANGPTL4, CA2, HAS3, HBEGF, KRT15, NRIP1, P2RY2, and SQSTM1) showed an increase in expression level by the combinations of 0.1 μM or 0.5 μM RP with Nam and *CS* (Figure 2). In comparing to the other retinoids, 0.1 μM tRA shows an inconsistent pattern of higher expression when compared to 0.1 μM or 0.5 μM RP. Additionally, 0.1 μM ROL had overall lower induction levels compared with 0.1 μM RP. In contrast to RARα activation, RP with Nam did not show a consistent pattern of increased expression over RP alone on the 40 responsive genes (data not shown).

**Figure 2.**
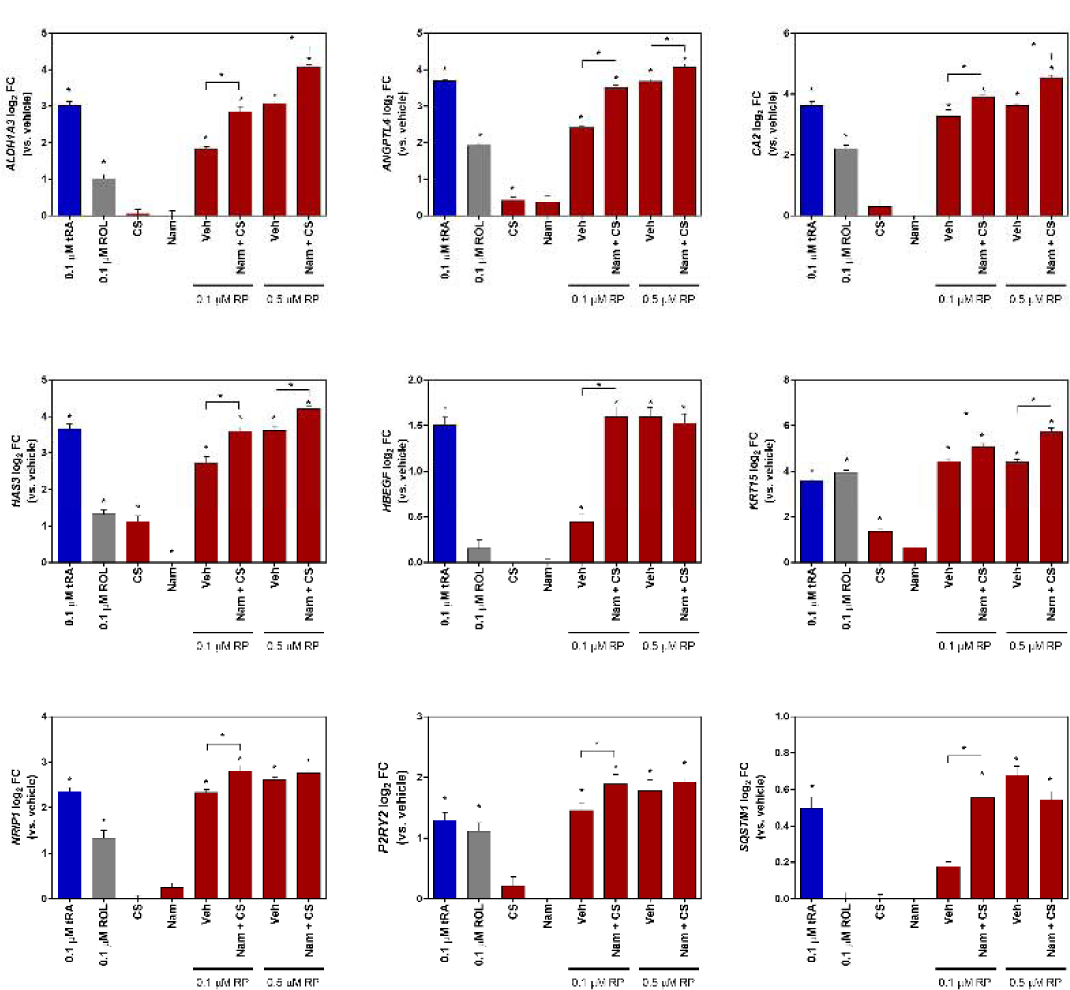
Retinoid responsive gene expression quantification shows increased response to combination of RP with niacinamide and *Ceratonia siliqua*. Quantitation of retinoid responsive target genes by qPCR in hTERT keratinocytes after treatment for 24 hours. Shown are the log2 fold changes (FC) of 9 select retinoid target genes (versus untreated). These include ALDH1A3, ANGPTL4, CA2, HAS3, HBEGF, KRT15, NRIP1, P2RY2, and SQSTM1. Treatments included single material of 0.1 μM tRA (blue bar), 0.1 μM ROL (gray bar), 1% *Ceratonia siliqua* (*CS)*, 1 mM niacinamide (Nam), 0.1 μM RP, or 0.5 μM RP (red bars). Additionally, a treatment combination of 0.1 μM RP or 0.5 μM RP with 1% *CS* and 1 mM Nam (red bars) was performed. Student’s t-test, \**p* < 0.05. n = 12 per treatment group. Error bars represent log2 SEM.

In conclusion, these collective data provide a body of evidence that RP in combination with Nam and *CS* can significantly increase RP activity in retinoid responsive *in vitro* models. Mechanistically, we believe this is via an increase in cellular NAD+ pools by the precursor Nam which allows for optimized oxidation to tRA via ROL and RAL. Additionally, the total flavonoids present in *CS* may further boost metabolism efficiency by inhibiting the reduction of RAL back to ROL (Figure 3). The *CS* carob fruit extract tested in this work also contains a high level of oligogalactomannans (data not shown, analysis by Silab, France). Thus, it is possible that these chemistries may also have a functional role in increasing the RAR response in the RP combinations. To better address these questions, future work is needed to confirm the retinoid metabolite profile from these retinoids and combinations in keratinocytes. We hypothesize that this combination has the potential to provide a stronger breadth and depth of efficacy response than RP alone and future work is needed to confirm this in clinical testing.

**Figure 3.**
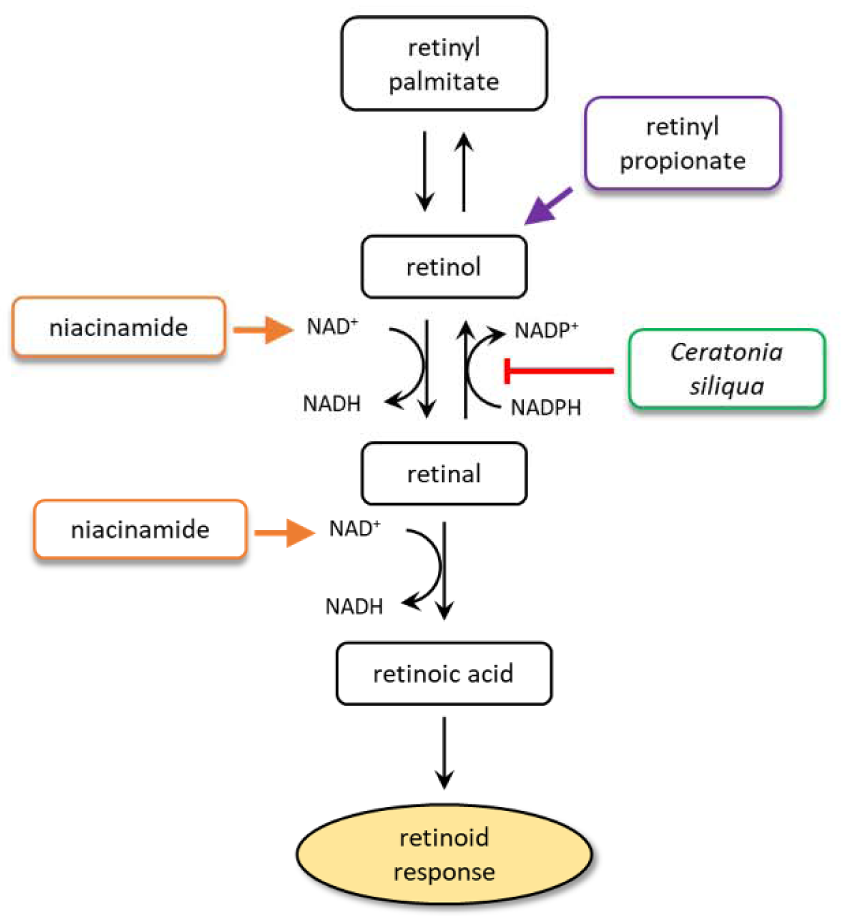
Schematic diagram of retinoid metabolism and hypothesized intervention points by niacinamide and *Ceratonia siliqua* for increased retinoid response when both are combined with RP. Enzymatic conversion of RP, RPalm, and ROL to tRA is well known and utilizes NAD^+^ as a key cofactor for the oxidation steps of ROL to RAL and RAL to tRA. Niacinamide, a known NAD+ precursor, is proposed to further heighten conversion by increasing NAD^+^ cellular levels. *Ceratonia siliqua (CS)* is proposed to inhibit the reverse reduction of RAL to ROL by functioning as a potent blend of flavonoids (104 μg/ml).

## Abbreviation List

RP: retinyl propionate
Nam: niacinamide
*CS*: *Ceratonia siliqua*
tRA: *trans*-retinoic acid

## ACKNOWLEDGEMENTS

We thank Holly Rovito and John Bierman for assistance with keratinocyte qPCR experiments.

